# Spreading depolarizations exhaust neuronal ATP in a model of cerebral ischemia

**DOI:** 10.1101/2024.07.30.605834

**Authors:** Karl Schoknecht, Felipe Baeza-Lehnert, Johannes Hirrlinger, Jens P. Dreier, Jens Eilers

## Abstract

Spreading depolarizations (SDs) have been identified in various brain pathologies. SDs increase the cerebral energy demand and, concomitantly, oxygen consumption, which indicates enhanced synthesis of adenosine triphosphate (ATP) by oxidative phosphorylation. Therefore, SDs are considered particularly detrimental during reduced supply of oxygen and glucose. However, measurements of intracellular neuronal ATP ([ATP]_i_), ultimately reporting the balance of ATP synthesis and consumption during SDs, have not yet been conducted.

In this study, we investigated neuronal ATP homeostasis during SDs using 2-photon imaging in acute brain slices from adult mice, expressing the ATP sensor ATeam1.03^YEMK^ in neurons. SDs were induced by application of potassium chloride or by oxygen and glucose deprivation (OGD) and were detected by recording the local field potential, extracellular potassium, as well as the intrinsic optical signal.

We found that, in the presence of oxygen and glucose, SDs were accompanied by a substantial but transient drop in neuronal [ATP]_i_. OGD, which prior to SD was accompanied by a slight reduction in [ATP]_i_ only, led to an even larger, terminal drop in [ATP]_i_ during SDs. Subsequently, we investigated whether neurons could still regenerate ATP if oxygen and glucose were promptly resupplied following SD detection. The data show that ATP depletion was essentially reversible in most cells.

Our findings indicate that SDs are accompanied by a substantial increase in ATP consumption beyond production. This, under conditions that mimic reduced blood supply, leads to a breakdown of [ATP]_i_. Therefore, our findings support therapeutic strategies targeting SDs after cerebral ischemia.

**Significance statement:** Spreading depolarizations occur in many brain pathologies, and are suggested to contribute to brain damage through a mismatch in energy supply and demand. However, measurements directly demonstrating an imbalance between ATP production and consumption, particularly in individual neurons, have not been reported. Here we show that spreading depolarizations lead to a transient decrease in intracellular neuronal ATP even in presence of ample glucose and oxygen. When the supply of oxygen and glucose was interrupted, ATP declined gradually until spreading depolarization led to the exhaustion of neuronal ATP. This process would be terminal without the renewed supply of oxygen and glucose. Therefore, therapies targeting spreading depolarizations could preserve neuronal ATP or delay its loss in brain pathologies.

## Introduction

Spreading depolarizations (SD) have been electrocorticographically recorded with subdural electrodes during neurocritical care in various situations: during the dying process after cardiac arrest, during the development of brain death with continued systemic circulation, during the evolution of delayed ischemic infarcts after subarachnoid hemorrhage (SAH), following malignant hemispheric stroke (MHS) due to middle cerebral artery occlusion, after traumatic brain injury (TBI), in Moyamoya vasculopathy, and an increasing number of further conditions associated with a disturbance of brain energy metabolism (Carlson et al., 2019; Dömer et al., 2024; Dreier et al., 2019; Dreier, Major, et al., 2018; Hartings et al., 2020; Lückl et al., 2018; Woitzik et al., 2013). Two clinical trials of larger scale in SAH and TBI have found that SDs are associated with worse patient outcome (Dreier et al., 2022; Hartings et al., 2020). A recent study in MHS patients has shown that SD-induced depression of neuronal activity is an electrocorticographic indicator of infarct growth (Kowoll et al., 2024), consistent with findings from animal studies indicating that interventions antagonizing SDs reduce cerebral damage (MacLean et al., 2023; Schoknecht et al., 2020; Yi et al., 2019). The SD continuum describes the spectrum from transient waves with negative direct current (DC) shifts of short to intermediate duration in adequately supplied or less ischemic tissue, to terminal waves in severely ischemic tissue characterized by long-lasting DC shifts and transition of the neurons from the state of injury to cell death (Dreier & Reiffurth, 2015; Hartings et al., 2017). Thus, SD recordings are indicative of the neuronal metabolic state.

SDs are characterized by a near-complete depolarization of neurons and glial cells, and a near complete breakdown of transmembrane ion gradients accompanied by a net influx of cations and water into the neurons (Dreier, 2011). From an electrochemical and thermodynamic point of view, SD is the abrupt, passive transition from a state of low entropy to a state of high entropy, which involves the loss of energy stored in electrochemical gradients (Dreier et al., 2013; Lemale et al., 2022). Subsequently, the recovery of cells from SD, i.e. the restoration of ion gradients, requires chemical energy, e.g. adenosine triphosphate (ATP). There is compelling evidence that activation of Na^+^/K^+^-ATPases is paramount to terminate SD and prevent neuronal death due to prolonged intracellular overload, e.g., with Na^+^ (Geisberger et al., 2021; Hernansanz-Agustín et al., 2020) and Ca^2+^ (G. Somjen, 2004). Increases in the intracellular concentration of sodium ions ([Na^+^]_i_) from ∼10 to ∼35 mM and in the extracellular concentration of potassium ions ([K^+^]_o_) from ∼3 to ∼10 mM, concentrations which are reached or even exceed during SDs, were shown to strongly stimulate Na^+^/K^+^-ATPases (Crambert et al., 2000; Horisberger & Kharoubi-Hess, 2002; Larsen et al., 2014). This causes a rise in ATP consumption and demand. In response, ATP production by oxidative phosphorylation is thought to increase, which has been experimentally demonstrated by measurements of higher cerebral metabolic rate of oxygen (CMRO_2_) (Mayevsky & Weiss, 1991; Piilgaard & Lauritzen, 2009; Schoknecht, Maechler, et al., 2023). At the same time, overall tissue ATP concentrations were shown to decrease by around 50% even in otherwise healthy parenchyma despite increased ATP synthesis in some studies (Mies & Paschen, 1984; Selman et al., 2004). These data support the hypothesis of reduced neuronal intracellular ATP concentrations ([ATP]_i_) during SDs, however, corresponding measurements, particularly at single-cell resolution, have not yet been conducted.

Here, we monitored neuronal [ATP]_i_ during SDs in acute brain slices from mice expressing the Förster resonance energy transfer (FRET) based ATP sensor ATeam1.03^YEMK^ under the neuronal promotor Thy1.2 (Imamura et al., 2009; Trevisiol et al., 2017). We monitored, to our knowledge for the first time, neuronal [ATP]_i_ during SDs in situ, first in the presence of oxygen and glucose, then during oxygen and glucose deprivation (OGD), and finally upon the renewed supply of oxygen and glucose after real-time detection of SDs.

## Results

### Imaging of [ATP]_i_ dynamics during SDs

To monitor neuronal [ATP]_i_, we acquired time series of the neuronally expressed ATP sensor (Fig. 1A) (Imamura et al., 2009; Trevisiol et al., 2017), either in multiple brain regions (referred to as ‘large scale’ imaging) or in up to a dozen individual neurons at single-cell resolution. The ratio of YFP over CFP of the FRET-based sensor provided a semi-quantitative readout of [ATP]_i_, hereafter referred to as ‘ATP signals’. In combination with established methods to induce SDs, i.e. by application of KCl (3M) (Fig. 1A ‘KCl puff’) or OGD, and to detected SDs, i.e. by electrophysiological recordings of the direct current (DC) local field potential (LFP), of the extracellular potassium ion concentration ([K^+^]_o_), and of the intrinsic optical signal (IOS, Fig. 1A) (Müller & Somjen, 1999; Schoknecht et al., 2020; Schoknecht, Hirrlinger, et al., 2023), we were able to investigate neuronal [ATP]_i_ in the course of SDs.

**Fig. 1.**
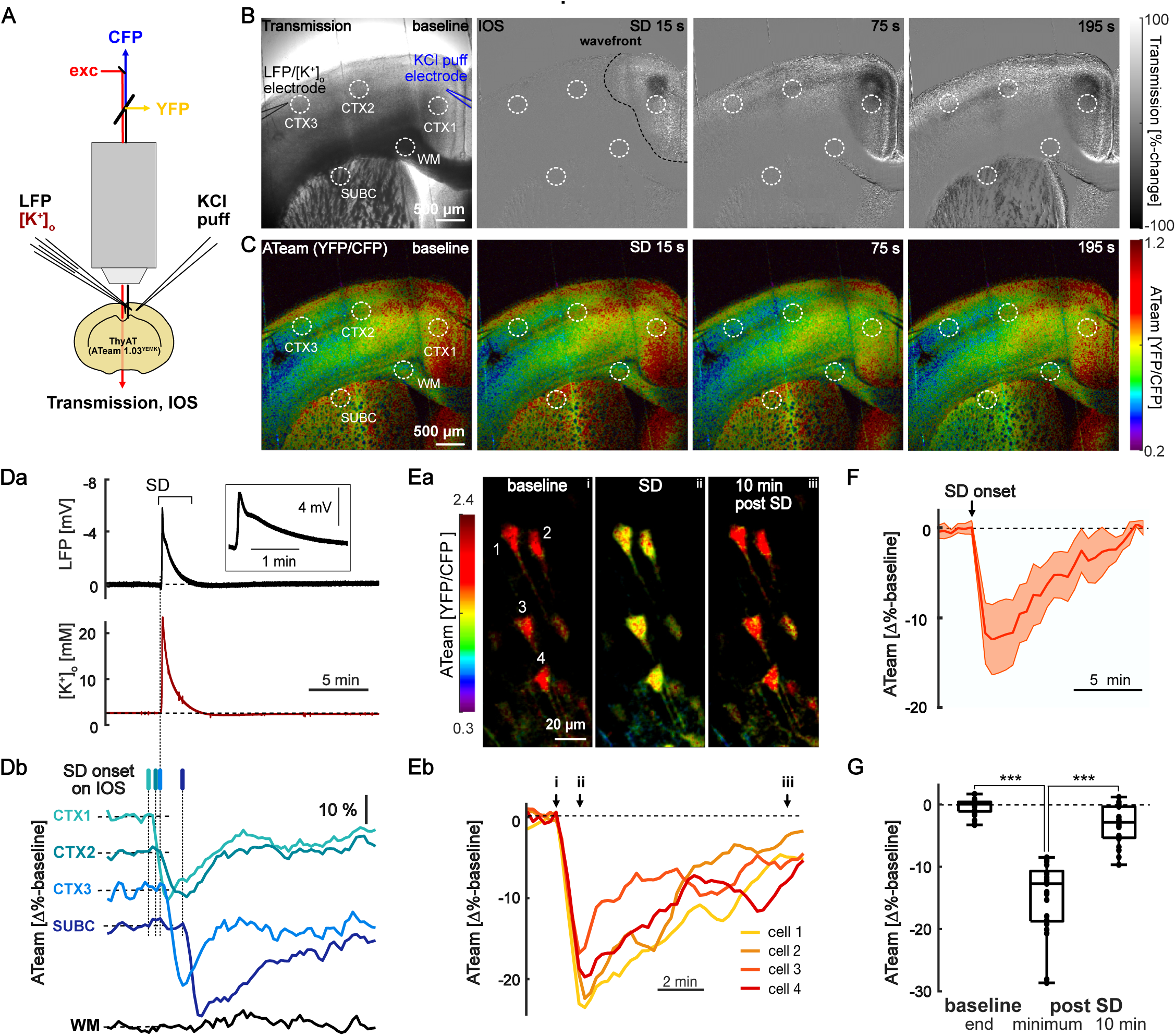
Spreading depolarizations induce a transient drop in neuronal ATP. A) Experimental setup. Spreading depolarizations (SDs) were evoked by puff application of KCl (3 M, right), while the local field potential (‘LFP’) and the extracellular K^+^ concentration (‘[K^+^]_o_’) were recorded by double-barreled electrodes (left) in acute mouse brain slices. Laser-scanning microscopy with two-photon excitation (‘exc’) was used to record the fluorescence of the ATP-sensor ATeam 1.03^YEMK^ (expressed under the neuronal promoter Thy1.2) as well as the transmission and the intrinsic optical signal (‘IOS’, i.e., the %-change in transmission). The Förster energy transfer (FRET) signal, i.e. the ratio of the acceptor and donor fluorescence (from yellow and cyan fluorescence proteins, ‘YFP’ and ‘CFP’, respectively), correlates to the intracellular ATP concentration ([ATP]_i_). B) Left: Transmission image of an acute mouse brain slice (300 µm thickness) prior to induction of an SD (‘baseline‘) by K^+^ puff application (1s). Black and blue lines outline the combined local field potential/extracellular potassium electrode (LFP/[K^+^]_o_) and the KCl puff electrode, respectively. Dotted circles denote regions of interest (ROIs), three in the neocortex, one subcortical and one in the white matter (CTX1-3, SUBC, and WM, respectively). Middle and right: IOS images 15, 75, and 195 s after SD induction. The dashed line in the ’15 s’ IOS image denotes the wavefront of the SD. Note that the SD propagates from the induction site to neocortical and subcortical regions, yet leave out the WM. Note how the SD propagates from the induction site (‘KCl puff electrode’ near CTX1) to the other neocortical and subcortical ROIs, yet leave out the WM. C) Corresponding color-coded images of the ATeam signal (YFP/CFP) based on the maximum intensity z-projection (7 z-planes at 5 µm intervals) of ratio images of the donor (CFP) and the acceptor-domain (YFP). From left to right: ATeam-signal before (‘baseline’) and after SD induction. Note how the transient reduction of the ATeam signal follows the SD (cf. B) first reaching the cortical (75s) ROIs, then the subcortical ROI (195s). Cooler colors indicate lower [ATP]_i_. D) a) Parallel LFP (top) and [K^+^]_o_ signals (bottom), corresponding to the experiment shown in B and C. Note the transient negative shift of the direct current (DC) potential in the LFP signal as well as the transient increase in [K^+^]_o_ (bottom), characteristic for SDs. Also note that the LFP trace is oriented according to the convention of electroencephalography (EEG) with negativity up and positivity down. Inset: Temporally enlarged, M-shaped LFP signal typical for SDs. The dotted vertical line continues to Db illustrating the relative timing of the shown signals. b) ATP signals from the ROIs indicated in B and C during the SD. Note the propagation of the SD from ‘CTX1’ to ‘SUBC’ and that the WM was not affected by the SD. The vertical bars indicate the onset of SD as detected by the IOS per ROI. E) a) Single-cell resolution, color-coded ratio images (20 z-planes,1.5 µm interval) capturing 4 neurons (labeled 1-4) before (left), during (middles) and 10 minutes after an SD (right) induced by K-puff application. Letters in top right corner (i, ii, iii) mark the timepoint of imaging corresponding to the curves shown in Eb. b) Self-normalized temporal ATeam profiles of the 4 neurons labeled in Da. Note the synchronized reduction followed by recovery of [ATP]_i_ in all 4 neighboring neurons. F) Average of ATeam-profiles synchronized to the SD onset from 99 self-normalized cells captured during 21 SDs (median ± IQR/2, 14 slices, 8 mice). G) Summary boxplot of ATeam-signals shown in E and F at the end of the baseline normalized to the baseline mean (typically 3-5 minutes, but up to ∼1 hr in a subset of slices (see Suppl. Fig. 2A), at the SD peak, and 10 min after the SD. Statistics are based on mean values of 2 to 12 cells captured per ROI during individual SDs (n=21 SDs, 14 slices, 8 mice, ***-p<0.001, Wilcoxon-test & Bonferroni correction). Box plots indicate the median, IQ1, and IQ3, whiskers the extrema; individual data points are indicated by dots.

### SDs induce a transient drop in neuronal ATP

We first addressed the question of whether ATP synthesis matches demand during SDs in neurons, here in presence of oxygen and glucose. Focal induction of SDs by KCl (Fig. 1A,B) led to SDs propagating across the neocortex to subcortical regions, as indicated by changes in the IOS, reflecting changes in water distribution between the intra- and extracellular space, which affects refractive tissue properties (Fig. 1B) (Müller & Somjen, 1999; Schoknecht, Hirrlinger, et al., 2023). The DC-potential showed a negative shift and [K^+^]_o_ increased sharply upon SD onset at the site of the recording electrode (Fig. 1Da). Parallel large-scale imaging of [ATP]_i_ revealed a substantial but transient reduction of the ATP sensor signal by 8-16% in cortical as well as subcortical regions of interest (ROIs, Fig. 1C,Db). Notably, the changes in [ATP]_i_ follow the propagation pattern of the SD (Fig. 1C,Db, cf. 1B). Neither IOS nor [ATP]_i_ changed in the white matter ROI, consistent with the finding that SDs are a gray matter phenomenon (Dreier, Lemale, et al., 2018).

Imaging at single-cell resolution confirmed the transient reduction in [ATP]_i_ during SDs in individual neurons (Fig. 1Ea,Eb). On average, the neuronal ATP signal decreased by 13% [10%, 18%] (median, IQ1 and IQ3), but recovered in all experiments (Fig. 1F,G, n=21 SDs, 14 slices, 8 mice, 99 individual neurons; p<0.001, Wilcoxon signed rank & Bonferroni correction). In summary, neuronal [ATP]_i_ levels drop transiently during SDs, even when oxygen and glucose are available.

To assure long-term stability of the ATP signal, we performed recordings for ∼55 min without experimental intervention (n=17 cells, 3 slices, 3 mice, Suppl. Fig. 1 A), revealing stable neuronal [ATP]_i_. Furthermore, to control for SD-induced neuronal swelling (Kirov et al., 2020; Zhou et al., 2010), we performed recordings in which a 30% hypoosmolar solution was applied (n=20 cells, 4 slices, 2 mice, Suppl. Fig. 1B,C). The hypoosmolar challenge, known to induce i.a. neuronal swelling (Murphy et al., 2017), was accompanied by a slight increase in the ATP signal, indicating that SD-induced drops in ATP signals (cf. Fig. 1) could be slightly underestimated.

### SDs deplete neuronal ATP during OGD

SDs were shown to occur in conditions of primarily reduced energy supply but could as well reduce [ATP]_i_ by increasing energy consumption (Ayata & Lauritzen, 2015; Lemale et al., 2022; Lindquist, 2024). This led us to investigate the effect of SDs on [ATP]_i_ in the context of cerebral ischemia, here OGD.

OGD was induced by wash in of artificial cerebrospinal fluid (aCSF), wherein glucose was replaced by sucrose and oxygen displaced by nitrogen. Recordings of partial tissue oxygen pressure (pO_2_) during wash in of the OGD-modified aCSF confirmed a critical reduction of the pO_2_ below the hypoxia threshold (∼8-10 mmHg, Fig. 2A) within approximately one minute (n=5 experiments), a pO_2_ level, at which oxidative phosphorylation was shown to break down (Kasischke et al., 2011).

**Fig. 2.**
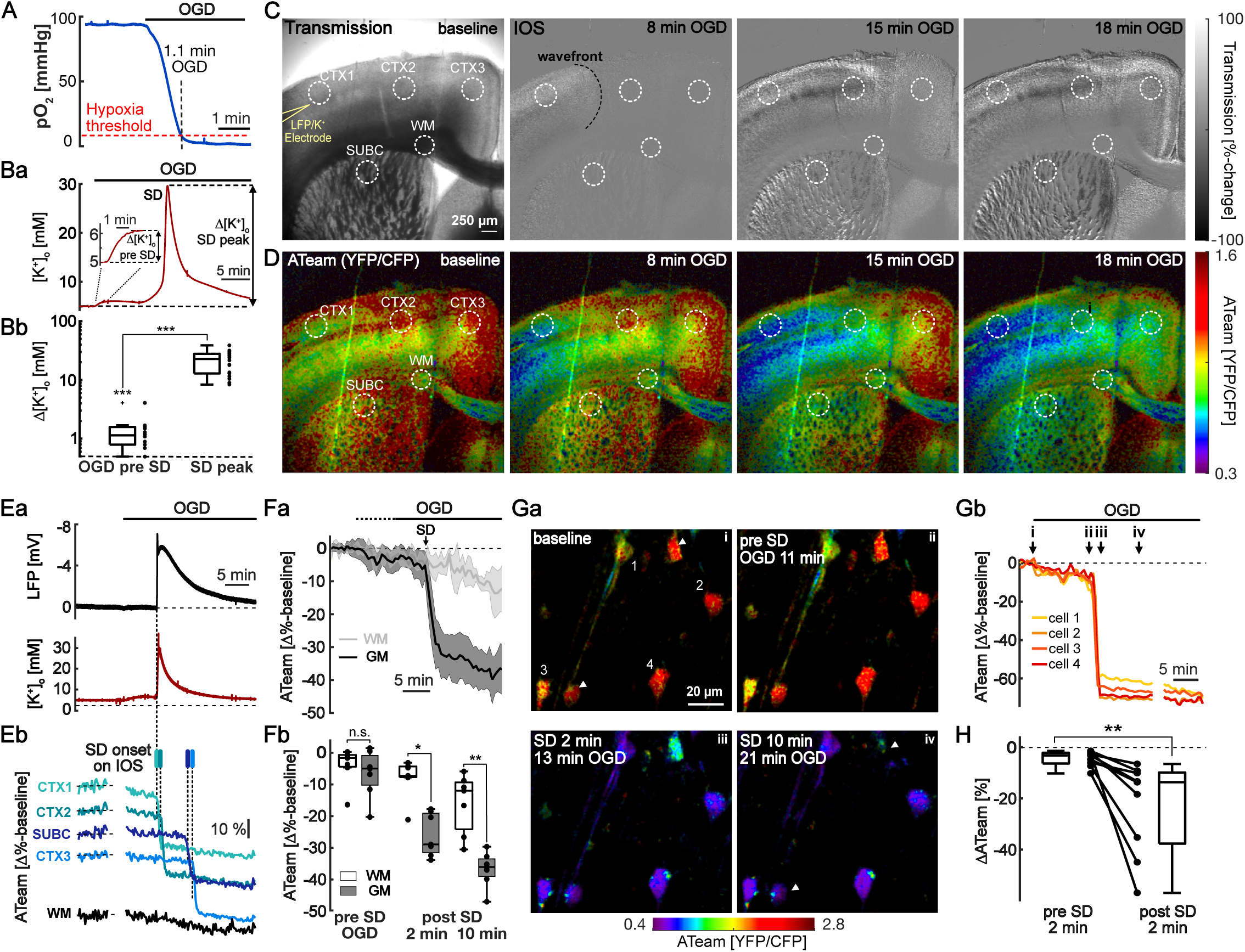
SDs deplete neuronal ATP during oxygen-glucose deprivation. A) Clark-electrode recording of partial oxygen pressure (pO_2_) 50 µm below the slice surface during wash-in of glucose-free and oxygen-depleted aCSF (OGD). Note, that the hypoxia threshold for the breakdown of oxidative metabolism (Kasischke et al., 2011) was reached in just over a minute. B) a) Recording of [K^+^]_o_ during wash-in of OGD solution, which shows a typical small early increase in [K^+^]_o_ (‘τι[K^+^]_o_ pre SD’) followed by a plateau. The occurrence of an SD was accompanied by a sharp rise in [K^+^]_o_ (τι[K^+^]_o_ peak). Note the delayed recovery of [K^+^]_o_ following the SD (cf. Fig. 1Da). b) Summary boxplot of the corresponding [K^+^]_o_ parameters (n=18 SDs in 18 slices, 11 mice; ***-p<0.001, Wilcoxon-test & Bonferroni correction). Box plots indicate the median, IQ1, and IQ3, whiskers the extrema; individual data points are indicated by dots. C) Transmission image (left) and IOS maps (from left to right) captured after the onset of a spontaneously occurring SD in ODG. The black dotted line in the IOS map ‘8 min of OGD’ indicates the wavefront of SD. D) Color-coded images of [ATP]_i_ synchronous to the IOS images (C) of the SD-associated reduction of [ATP]_i_ in OGD. E) a) LFP- and [K^+^]_o_-recording corresponding to the experiment in C and D, showing the prolonged SD (top) occurring in OGD (cf. Fig. 1Da) as well as a subtle increase in [K^+^]_o_ early during OGD and the prolonged increase in [K^+^]_o_ during the SD. b) ATP signals from the ROIs indicated in C and D during the SD in OGD. The vertical bars indicate the onset of SD as detected by the IOS per ROI. Note that the sharp decline in [ATP]_i_ coincides with the onset of the SD. F) a) Average of ATeam signal in gray and white matter ROIs (‘GM’ and ‘WM’, respectively); synchronized to the SD onset in each ROI (median ± IQR/2; 28 GM ROIs, 7 WM ROIs, 7 slices, 5 mice). Note that SDs started at variable times in OGD (indicated by dotted line on top), in this subset of slices after 9.8 min [7.2 min, 12.6 min] (median, IQ1 and IQ3). b) Boxplots summarizing the self-normalized ATeam signals (as shown in Fa) in OGD prior to SDs and 2 and 10 minutes after SDs. Boxplot shows the median, IQ1, and IQ3, whiskers the extrema; individual data points are indicated by dots (n=7 slices, 5 mice, n.s.-not significant, *-p<0.05, **-p<0.01, Mann-Whitney U-test & Bonferroni correction). G) a) Single-cell resolution ATeam images showing individual neurons before OGD (top left, ‘baseline’), following 11 min in OGD (top right, ‘pre SD’), and after SD (bottom left and right). Arrowheads point to neurons that moved out of the focal planes, which were not analyzed. Letters in top right corner (i, ii, iii, iv) mark the timepoint of imaging corresponding to the curves shown in Gb. b) Self-normalized ATP signals of the 4 neurons labeled in Ga. Note that the [ATP]_i_ did not recover from the SD-induced drop. H) Boxplot summarizing the change in ATP signal in OGD during the 2 min before SD onset (left), and during the first 2 min after SD onset (right; n=9 slices, 6 mice, 45 individual neurons, Wilcoxon test, *-p<0.01). SDs started after 12.8 min [12.3 min, 15.4 min] of OGD.

As an early effect of OGD, [K^+^]_o_ increased on average by 1.2 mM [0.8 mM, 1.6 mM], (n=18 SDs in 18 slices, 11 mice; p<0.001, Wilcoxon-test & Bonferroni correction; Fig. 2B), typically reaching a plateau (Fig. 2Ba). SDs occurred spontaneously but at varying timepoints in OGD, on average after 12.9 min [12.0, 16.0] (median, IQ1 and IQ3; n=36 slices, 19 mice, 34 SD-positive slices) and were accompanied by sharp increments in [K^+^]_o_ by 22.9 mM [13.0 mM, 28.0 mM] (n=18 SDs in 18 slices, 11 mice; p<0.001, Wilcoxon-test & Bonferroni correction; Fig. 2B, Ea) and the typical negative DC-shift (Fig. 2Ea). SDs propagated across the cortex to subcortical gray matter as shown by the IOS, similar to KCl induced SDs (Fig. 2C).

[ATP]_i_ decreased gradually in OGD, followed by a substantial, sharp drop upon spontaneous SD onset, which in contrast to well-nourished slices, did not recover (Fig. 2D,Eb, cf. Fig.1). Before SD onset the ATP sensor signal declined to -5% [-1%, -10%] compared to baseline values before OGD over a time course of 9.8 min [7.2 min, 12.6 min] in regions that eventually underwent SD, i.e. the gray matter (n=7 slices, 5 mice, Fig. 2Fa-b). Notably, the ATP signal decreased further to -29% [-19%, -32%] and -36% [-33%, -39%] at two and 10 minutes after SD onset. The time points were chosen to capture the prompt SD-induced decline in [ATP]_i_ as wells as the absence of acute recovery. After SD, [ATP]_i_ levels were significantly lower in ROIs that underwent SDs compared with those that did not, i.e. in the white matter (n=7 slices, 5 mice, p>0.05, Mann-Whitney U-test, Fig. 2Fa-b). Single-cell imaging confirmed a sharp drop in neuronal [ATP]_i_ without recovery following SD (Fig. 2Ga-b,H). The ATP signal dropped by 3% [2%, 6%] in the 2 minutes prior to SD compared to a drop of 14% [10%, 38%] in the first 2 minutes after SD (Fig. 2H, n=9 slices, 6 mice, 45 cells in total, p<0.01, Wilcoxon-test). The single-cell data confirm that [ATP]_i_ did not recover in OGD.

### SD-induced ATP depletion under OGD is reversible

We next investigated whether neurons are able to restore [ATP]_i_ following the significant drop after SDs in OGD, or whether [ATP]_i_ depletion is terminal. For this purpose, we once more acquired [ATP]_i_ signals during a period of OGD (Fig. 3A), but this time started to reperfuse oxygen- and glucose-containing aCSF upon real-time detection of SDs in the electrophysiological recordings (Fig. 3Ba).

**Fig. 3.**
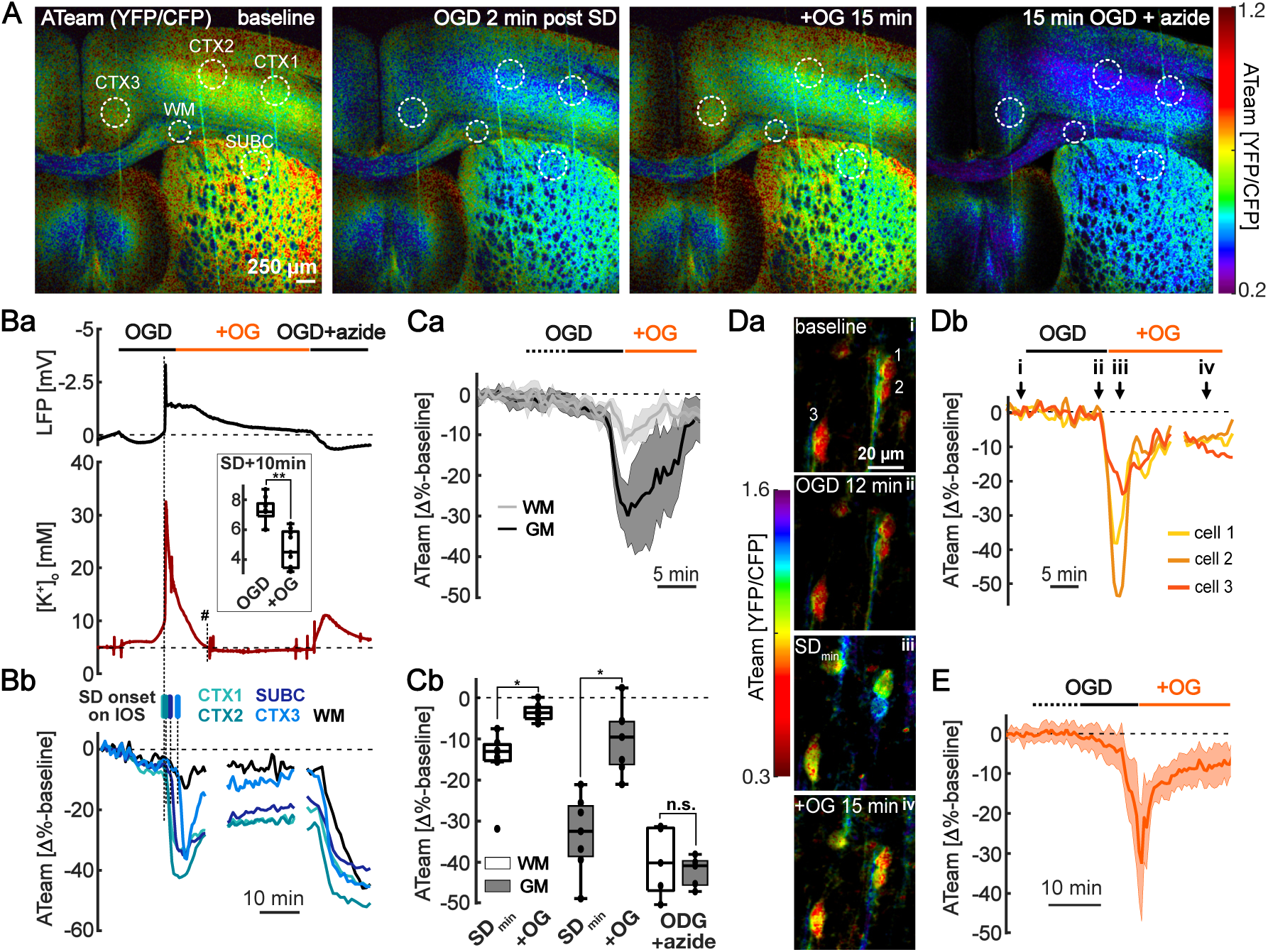
Reversibility of SD-induced ATP depletion during oxygen-glucose deprivation. A) From left to right: ATeam maps before OGD (‘baseline’), 2 min post SD in OGD, 15 minutes following re-supply of oxygen and glucose (‘+OG’) and 15 min after wash-in of OGD together with sodium azide (inhibitor of complex IV). White dotted circles mark three neocortical ROIs (CTX1-3), one subcortical (SUBC) and one white matter (WM) ROI, analyzed in Bb. B) a) LFP- and [K^+^]_o_-recording corresponding to A, confirming a spontaneously-occurring SD in OGD. Note the sharp decline in [K^+^]_o_ upon re-supply of OG, which slightly undershoots baseline levels. Vertical line and ‘#’ denotes timepoint 10 minutes after the [K^+^]_o_ peak during SD, which was used for quantitative analyses shown in inset. Brief, spike-like transients (for example those next to ‘#‘) are typical recording artifacts. Inset: Boxplot of [K^+^]_o_ (in mM) 10 minutes after its peak during SD in persistent OGD (left bar; data derived from experiments for Fig. 2 (see Ba,Ea)) and for re-supply of OG (right bar, n=9 slices each, 5 & 6 mice, **-p<0.01, Mann-Whitney U test). b) ATP signals from the ROIs indicated in A. The vertical bars indicate the onset of SD as detected by the IOS per ROI. Note that, [ATP]_i_ recovered upon re-supply of OG in all ROIs. Also note, that the addition of sodium azide during OGD lowered [ATP]_i_ in the WM similar to the ROIs from the GM. C) a) Average of transients in the ATeam signal separated into ROIs from GM and WM synchronized to the SD onset (median ± IQR/2; 28 GM ROIs, 7 WM, 7 slices, 5 mice). SD started 14 min [12.8 min, 15.8 min] after OGD in this subset of slices. b) Boxplots of the self-normalized ATeam signal (as shown in Ca) quantifying the minimum after SD in OGD (‘SD_min_’), 15 min after resupply of oxygen and glucose (‘+OG’), and in OGD together with sodium azide (SD_min_ and +OG; n=7 slices, 5 mice, *-p<0.05, Wilcoxon test & Bonferroni correction; OGD + azide, n=5 slices, 4 mice, Mann-Whitney U-test, p=0.63, n.s.-not significant). D) a) From top to bottom: ATeam maps showing individual neurons before OGD, 12 min in OGD (immediately before SD onset), during the SD (‘SD_min_’), and 15 min after re-supply of OG (‘+OG’). Letters in top right corner (i, ii, iii, iv) mark the timepoint of imaging corresponding to the curves shown in Db. b) Self-normalized transients of the ATeam signal of the 3 neurons labeled in Da. E) Average of ATeam transients of neurons that showed partial [ATP]_i_ recovery upon re-supply of OG (median ± IQR/2, n=11 slices, 8 mice; out of 60 neurons, 39 showed a recovery). Data are synchronized to the start of +OG. 21 of 60 cells showed no recovery of [ATP]_i_ (not shown). In this subset of experiments, SDs started 15.3 min [12.2 min, 18.4 min] after OGD.

These experiments resembled those shown in Fig. 2 up to the point at which oxygen and glucose were resupplied (‘+OG’), both at large-scale and single-cell imaging (Fig. 3A-C and Fig. 3D,E, respectively). Resupply of oxygen and glucose led to a return of the negative shift in the DC-potential (Fig. 3Ba) and a full restoration of [K^+^]_o_ to baseline levels with a transient undershoot (Fig. 3Ba). Ten minutes after the SD-associated peak of [K^+^]_o_, [K^+^]_o_ had returned to baseline levels or even below (4.5 mM [3.5 mM, 5.9]) in slices with renewed supply of oxygen and glucose, whereas [K^+^]_o_ remained elevated during persistent OGD (7.2 mM [6.9 mM, 7.8 mM]; Fig. 3Ba inset; n=9 slices from 5 & 6 mice, p<0.01, Mann-Whitney U test).

Notably, resupply of oxygen and glucose induced a prompt recovery of [ATP]_i_ (Fig. 3A, Bb). On average, ATP signals increased significantly towards baseline levels (baseline corresponds to 0% change), namely from -13% [-11%, -15% ] to -4% [-2%, -5%] and from -32% [-26%, -38%] to -10% [-6%, -16%] in white matter ROIs and gray matter ROIs, respectively, following the resupply of oxygen and glucose for a period of 15 min (Fig. 3C, n=7 slices, 5 mice, p<0.05; Wilcoxon test & Bonferroni correction). To test whether the differences in ATP signals in SD-affected areas compared to unaffected areas were due to different basal ATP levels (Köhler et al., 2020), we performed a one-point calibration of the ATP sensor by applying sodium azide in addition to OGD (inhibitor of complex IV). Sodium azide has been shown to promptly deplete neuronal ATP (Pape & Rose, 2023; Trevisiol et al., 2017). Consistently, we saw a sharp reduction in [ATP]_i_ to similar levels independent of brain regions upon wash-in of sodium azide (Fig. 3Bb, Cb; Mann-Whitney U-test, p=0.63, n=5 slices, 4 mice), indicating a similar range of the ATP sensor signal and basal [ATP]_i_ in all ROIs. Lastly, we investigated recovery of [ATP]_i_ following SDs at the cellular level and found that about two thirds of the neurons were essentially capable to restore [ATP]_i_ (Fig. 3 D,E). Taken together, brain tissue affected by SDs undergoes a critical, yet in principle reversible, reduction in neuronal [ATP]_i_ during OGD.

## Discussion

SDs have been suggested to contribute to developing brain injury in various brain pathologies, particularly when involving a mismatch in the supply and demand of energy ((Lemale et al., 2022)). Therefore, we investigated the effect of SDs on neuronal [ATP]_i_. We provide evidence that SDs increase neuronal ATP consumption beyond production, exhausting neuronal [ATP]_i_ during OGD, which could be restored by renewed supply of oxygen and glucose upon detection of SDs in most cells.

### Increased ATP consumption and production during SDs

There is evidence that both ATP consumption and ATP production increase following SDs, the former via stimulation of Na^+^/K^+^-ATPase activity due to rising [K^+^]_o_ and [Na^+^]_i_ ((Crambert et al., 2000; Horisberger & Kharoubi-Hess, 2002; Larsen et al., 2014) and Introduction) and the latter via stimulation of glycolysis and oxidative phosphorylation, as concluded from measurements of reduced brain glucose concentrations as well as increased cerebral metabolic rate of oxygen (CMRO_2_) ((Balança et al., 2017; Lourenço et al., 2017; Mayevsky & Weiss, 1991; Piilgaard & Lauritzen, 2009; Schoknecht, Maechler, et al., 2023)). Here, [ATP]_i_ was transiently reduced following SDs in presence of oxygen and glucose, which indicates ATP consumption in excess of production (Fig. 1).

Transient reductions in [ATP]_i_ upon putatively increased consumption during SDs could be explained by either delayed or exhausted adaptation of ATP production. It remains unclear whether ATP production is fully exploited in the course of SDs, however, there is evidence that adaptation of ATP production is indeed delayed. Previous experiments have shown that peak CMRO_2_, indicating stimulation of oxidative phosphorylation, follows SD onset by ∼30 seconds (Schoknecht et al., 2023). In addition, in primary neuronal cultures, ATP production by glycolysis and oxidative phosphorylation was coupled to [Na^+^]_i_-dependent stimulation of Na^+^/K^+^-ATPase activity and similarly delayed by several seconds (Baeza-Lehnert et al., 2019). In hippocampal slices, neuronal activation, accompanied by a Na^+^ influx (Gerkau et al., 2019), has been shown to induce a prolonged yet delayed increase in neuronal glycolysis (Díaz-García et al., 2017). Taken together, a delayed activation of glycolysis and oxidative phosphorylation, could explain a transient reduction of neuronal [ATP]_i_. During SDs, elevated [Na^+^]_i_ (stimulating the Na^+^/K^+^-ATPase) results from the sustained influx of Na^+^ through ionotropic NMDA-receptors, which are activated by glutamate (Mei et al., 2020). Extracellular glutamate was shown to reach levels of up to ∼100 µM during SDs (Menyhárt et al., 2022; Zhou et al., 2013). Notably, glutamate applied at concentrations of 50 µM and 1 mM led to a reduction of [ATP]_i_ in primary neuronal cultures and in acute hippocampal brain slices, respectively (Baeza-Lehnert et al., 2019; Gerkau et al., 2019). Direct application of NMDA (30 µM) similarly reduced [ATP]_i_ of cultured neurons, overall supporting Na^+^ influx via NMDA-receptors triggering Na^+^/K^+^-ATPase activity and ATP consumption, to be followed by ATP production (Baeza-Lehnert et al., 2019). In addition, reversible fragmentation of mitochondria, as observed during SDs in otherwise healthy animals *in vivo,* could delay ATP production and thus contribute to a transient neuronal [ATP]_i_ reduction (Sword et al., 2024).

Previous studies demonstrated high sensitivity of ATeam1.03^YEMK^ to [ATP]_i_ over ATP-independent effects in various conditions. In particular, following the application of NMDA, glutamate, as well as increased [K^+^]_o_, a mutated version of the ATP sensor with a dysfunctional ATP-binding site showed little or no signal changes compared to the functional sensor (Köhler et al., 2020; Lange et al., 2015), indicating that ATP sensor signals here and in previous studies reflect an actual decreases in [ATP]_i_ rather than an unspecific effect on the fluorophores of the ATP sensor.

[K^+^]_o_ recordings have provided evidence for increased ATP consumption due to increased Na^+^/K^+^-ATPase activity during SDs (Major et al., 2017; Schoknecht et al., 2023) and were confirmed here: first, by the rapid return of [K^+^]_o_ from its peak to baseline levels, and second, by the reduction in [K^+^]_o_ below baseline levels for several minutes after SD, also called ‘undershoot’ in [K^+^]_o_ (Haglund & Schwartzkroin, 1990; G. G. Somjen, 2001)). Notably, [ATP]_i_ began to recover despite ongoing, increased consumption by the Na^+^/K^+^-ATPase, indicating that ATP production rapidly adopts to and even increases beyond consumption. To allow gradual recovery of [ATP]_i_, Na^+^/K^+^- ATPase activity, while shown to activate glycolysis and oxidative phosphorylation (Baeza-Lehnert et al., 2019; Meyer et al., 2022), could in addition be regulated to operate, i.e. consume ATP, at rates slightly below ATP production. Consistently, experiments on inside-out vesicles indicated an inhibitory feedback-loop on Na^+^/K^+^- ATPase activity, showing reduced Na^+^ transport by the Na^+^/K^+^-ATPase when ADP levels were increased (Mercer & Dunham, 1981). ADP levels are expected to increase due to ATP-hydrolysis resulting from aforementioned stimulation of Na^+^/K^+^-ATPase activity in the course of SDs.

At the end of SDs, marked by normalization of the DC-potential, neuronal [ATP]_i_ was still reduced compared to baseline, but gradually increased further thereafter. A similar time course has been observed when overall brain ATP content was measured spectrophotometrically in lysates of brain samples collected at different timepoints after SDs from rats (Mies & Paschen, 1984), suggesting that these measurements were in principle representative of neuronal [ATP]_i_ dynamics in presence of glucose and oxygen. However, not all studies measuring ATP in tissue samples found reduced ATP after SDs (reviewed in (Ayata & Lauritzen, 2015)). We thus provide further and first cell-specific evidence, for reductions in neuronal [ATP]_i_ in the course of SDs.

### SDs exhaust neuronal ATP in affected gray matter

Given that SDs were observed in conditions of metabolic compromise while simultaneously increasing ATP consumption (Lindquist, 2024), we monitored [ATP]_i_ during OGD, focusing particularly on the effect of SDs on neuronal [ATP]_i_.

Remarkably, SDs occurred during OGD when neurons still contained considerable amounts of [ATP]_i_, which, in some cases hardly differed from pre OGD levels (Fig. 3Db). However, a sharp drop in [ATP]_i_ followed the SD onset. Thus, SDs were probably not the consequence of widespread depletion of neuronal [ATP]_i_, although we cannot exclude prior depletion of [ATP]_i_ at the very focal SD initiation site. [ATP]_i_ depletion during continuous OGD was terminal. Consistently, in rats with middle cerebral artery occlusion, ATP remained lower in tissue samples of the penumbra after spontaneous SDs than in tissue samples from non-ischemic animals after induced SDs (Selman et al., 2004). Prevention of SDs could protect a majority of neurons from both the loss of residual [ATP]_i_ as well as from the ATP demand necessary to fully restore transmembrane ion gradients. Based on our data, brain regions, which are more resistant to SDs or inherently do not undergo SD, such as white matter, are expected to retain ATP longer during OGD.

During OGD or ischemia in vivo, intracellular neuronal pH has been shown to decrease to 6.7-7.0 (Fujiwara et al., 1992; Kelmanson et al., 2021; Knöpfel et al., 1998; Melzian et al., 1996). In this pH range, the signal of the ATP sensor ATeam1.03^YEMK^, i.e. the YFP/CFP ratio, is partially pH-sensitive, however, predominantly for ATP concentrations above 1 mM. Below 1 mM, ATP signals were mostly pH-insensitive or would even increase slightly with acidification (Natsubori et al., 2020), i.e. opposite to what we observed. Neuronal [ATP]_i_ was estimated to be 2.0-2.8 mM (Pape & Rose, 2023; Pathak et al., 2015; Rangaraju et al., 2014). For [ATP]_i_ in this range, intracellular acidification to pH 6.7 could explain decreased ATP signals by 10-15%, but not by 70%, as seen in individual neurons (Fig. 2Gb), and, importantly, only if ATP would not drop in parallel, as this would result in a shift into the pH-insensitive range of the sensor (Natsubori et al., 2020). Furthermore, intracellular acidification during ischemia or OGD was shown to start immediately (Fujiwara et al., 1992; Kelmanson et al., 2021; Knöpfel et al., 1998; Melzian et al., 1996), whereas we detected the major drop in ATP in association with SDs, on average ∼13 min in OGD. Importantly, recovery of [ATP]_i_ due to the resupply of oxygen and glucose (Fig. 3), i.e. increasing ATP signals from low [ATP]_i_ cannot be explained by pH-normalization as the sensor would not respond to changes in pH but to changes in [ATP]_i_ for [ATP]_i_ of 0.5 mM or lower. SDs, in presence of oxygen and glucose, have also been reported to undergo a transient intracellular acidification, however only semi-quantitively (Zhou et al., 2010), which prevents an estimation on the contribution of the changes in pH to the observed ATP signal. Notably, the changes in pH were limited to the period of depolarization, whereas we detected a reduction in [ATP]_i_ beyond the SD duration. In summary, although we cannot exclude that some changes in ATP-sensor signal resulted from changes in pH, the signal kinetics, the magnitude (changes of up to 70%), and the recovery of [ATP]_i_ could not be dominated by changes in pH, suggesting that observed ATP sensor signals indeed report changes in [ATP]_i_.

In contrast to OGD, neuronal [ATP]_i_ decreased directly and sharply upon pharmacological inhibition of oxidative phosphorylation by sodium azide (Fig. 3), a finding previously described in organotypic neocortical slices (Pape & Rose, 2023) and considered to result from a direct neuronal influx of Na^+^ due to opening of TRPV4 (Pape & Rose, 2023). A Na^+^ influx would increase ATP consumption by activation of the Na^+^/K^+^-ATPase, while ATP synthesis would be reduced due to simultaneous inhibition of complex IV ((Baeza-Lehnert et al., 2019; Crambert et al., 2000; Horisberger & Kharoubi-Hess, 2002), cf. prior discussion). Notably, a mutated version of the ATeam sensor previously remained insensitive to the application of sodium azide (Köhler et al., 2020; Lange et al., 2015).

### Reversibility of neuronal ATP-depletion induced by SDs during OGD

To mimic reperfusion therapy in acute stroke patients, we resupplied oxygen and glucose upon detection of SD. Remarkably, [ATP]_i_ essentially recovered in all of the investigated ROIs and in about two thirds of neurons imaged at single-cell resolution (Fig. 3). *In vivo,* changes in [ATP]_i_ may follow different kinetics, e.g. due to blood flow responses, which can be impaired after stroke and thus limit the supply of oxygen and glucose during reperfusion (Dreier, 2011). Nevertheless, our data reveal that even after an anoxic SD, which would have been terminal without resupply of oxygen and glucose, mitochondria are functional or rapidly regain function. This opens up the question, which substrates fuel the renewed ATP synthesis, specifically whether extracellular lactate becomes a substrate of oxidative phosphorylation by neuronal uptake and conversion to pyruvate or whether accumulated intracellular lactate and renewed glycolysis play the predominant role. Lactate was shown to accumulate in the brain parenchyma after various conditions: following SDs and cerebral ischemia models *in vivo* and *in vitro* (Balança et al., 2017; Lourenço et al., 2017; Paschen et al., 1987), in patients following ischemic stroke (Woitzik et al., 2014), and, specifically in response to SDs, after traumatic brain injury, subarachnoid hemorrhage and cerebral hematoma (Feuerstein et al., 2010). Thus, lactate could in principle be used for oxidative phosphorylation if transported into neurons. Overall, the role of extracellular lactate as a substrate of neuronal oxidative phosphorylation remains debated (Bak & Walls, 2018; Barros & Weber, 2018; Kann, 2024; Rae et al., 2024). Direct measurements of [ATP]_i_, as shown in this study, could be used to identify cell types and substrates that allow neuronal recovery of [ATP]_i_ after SDs in transient OGD, with potential implications for stroke therapy.

In summary, we show that SDs are accompanied by a substantial increase in neuronal ATP consumption beyond production. Subsequently, SDs exhaust neuronal [ATP]_i_ under conditions that mimic reduced blood supply, whereas before SD onset, [ATP]_i_ decreased markedly slower. Still, most cell were capable to restore [ATP]_i_ if oxygen and glucose were rapidly resupplied following SD. Our findings support therapeutic strategies to prevent SDs after cerebral ischemia as this could prevent not only the acute loss of [ATP]_i_ but also the remaining need of energy to fully restore ion gradients. Future research may address the potency of therapeutics to preserve neuronal [ATP]_i_ by preventing SDs as well as the metabolic mechanisms that drive [ATP]_i_ recovery upon the renewed supply of oxygen and glucose.

## Material and Methods

### Animals

Animals were housed and bred in accordance with the German Animal Welfare Act, the European Communities Council Directive (2010/63/EU), as well as Leipzig University guidelines and with approval of the local authorities (T09/20, T05/21-MEZ). Animals had free access to food and water and were kept in a 12 h/12 h light-dark cycle. The study includes 12 female and 15 male adult transgenic mice (median age 23 weeks) expressing the ATP sensor ATeam 1.03^YEMK^ (Imamura et al., 2009) under the neuronal promotor Thy1.2 on the background of the C57BL/6J mouse strain (B6-Tg(Thy1.2-ATeam1.03^YEMK^)AJhi; MGI:5882597; referred to as ThyAT) (Trevisiol et al., 2017).

#### Preparation

Animals were anesthetized by isoflurane and decapitated. Brains were removed and cut into coronal slices (thickness 300 µm, -1.5 mm to 2 mm relative to bregma) on a vibratome (Leica VT1200S, Leica Biosystems, Nussloch, Germany) in ice-cooled artificial cerebrospinal fluid (aCSF), then transferred to warmed aCSF (35 °C) for 30 min and kept at room temperature (20-22°C).

#### Solutions and drugs

Drugs and chemicals were purchased from Sigma Aldrich. ACSF contained in mM: 125 NaCl, 2.5 or 5 KCl, 1.25 NaH_2_PO_4_, 26 NaHCO_3_, 1.8 MgCl_2_, 2 CaCl_2_, and 20 glucose and was continuously carbogenated (95% O_2_, 5% CO_2_, pH 7.4, osmolarity ∼305 mOsm). For oxygen-glucose deprivation (OGD), glucose was replaced by equimolar saccharose and gassed with a mixture of 95 % nitrogen (N_2_) and 5% CO_2_ (pH 7.4, osmolarity ∼305 mOsm). Complex IV of the respiratory chain was inhibited by sodium azide (5 mM) in a subset of experiments.

#### Electrophysiological recordings

For electrophysiological recordings and parallel imaging (see below), slices were transferred to an open bath perfusion chamber (Warner Instruments, Holliston, MA, USA, perfusion rate 6-8 ml/min). Slices were held in place with a U-shaped platinum wire covered with a nylon grid. Direct current (DC) local field potentials (LFP) were recorded ∼50 µm below the surface of the slice in layer 2-3 of somatosensory and - motor areas using microelectrodes prepared from borosilicate glass (Science Products, Hofheim, Germany; pipette puller PC-10, Narishige, Tokyo, Japan) or from double-barrel theta glass (Fa. H. Kugelstätter, Garching, Germany) to measure the LFP in parallel with K^+^-sensitive Nernst potentials. The potassium-selective barrel was filled with solution containing 150 mM KCl and Potassium-Ionophor 1 (Valinomycin) at the tip. The LFP and the Nernst potential were amplified (EXT-10 and ION-01 for LFP or LFP and [K^+^]_o_, respectively), low-pass filtered (3 kHz, LPBF-01GX, all npi electronic, Tamm, Germany) and digitized (CED-1401 Micro3 with Spike2 9.01 software, Cambridge Electronic Design, Cambridge). To confirm oxygen deprivation, partial tissue oxygen pressure (pO_2_) was recorded using Clark-style glass oxygen microelectrodes (10 µm tip; Unisense, Aarhus, Denmark) (Schoknecht et al., 2017) during wash in of OGD solution in a subset of experiments.

#### Induction of SD

SDs were either triggered focally by release of 3 M KCl from glass microelectrodes (2-3 MOhm when filed with ACSF) by pressurized air (0.5 bar, 1 s, PDES-2L, npi electronic, at >500 µm from the recording site) or occurred spontaneously during OGD. During OGD, SDs occurred in only 33% of the brain slices after 12.8-18.0 minutes (n=15, 5 mice, data not shown), a low incidence compared to *in vivo* models of cerebral ischemia, in which most animals developed SDs (see e.g. (Lückl et al., 2018; Schoknecht et al., 2020)). Given that [K^+^]_o_ is a trigger of SDs and is here constantly diluted by the perfused aCSF containing 2.5 mM K^+^, we modified the experimental approach by raising K^+^ to 5 mM in the aCSF. Importantly, at this concentration K^+^ by itself did not induce SDs (Reiffurth et al., 2019; Schoknecht, Hirrlinger, et al., 2023). Raising K^+^ in the solution increased the incidence of SDs to 94% (n=31 slices, 16 mice).

#### Intrinsic optical signal and FRET

Images were acquired by 2-photon laser scanning microscopy using a Ti:Sapphire laser (810 nm, Mai Tai DeepSee, Spectra-Physics, Milpitas, CA, USA) and a laser scanning microscope (Olympus Fluoview 10M, Olympus, Tokyo, Japan) equipped with a 3.32x (NA:0.14) or 25x objective (XLPlan N 25x W NA:1.05, both Olympus). We recorded time series of vertical stacks (image dimension 320×320 to 640×640 pixels, xy pixel dimension: 0.2-12 µm), up to 20 z-steps (1.5-5 µm/image plane) and time intervals of 15-40 seconds during SDs and up to 2 minutes for extended baseline measurements.

To detect onset and propagation of SDs, we measured light transmission as previously described (Schoknecht, Hirrlinger, et al., 2023). Refractive tissue properties, affect light transmission (and reflection), which shows characteristic changes during SDs and is also known as the intrinsic optical signal (IOS)(Müller & Somjen, 1999).

To measure neuronal ATP, we used the Förster resonance energy transfer (FRET)-based sensor ATeam 1.03^YEMK^ consisting of a donor domain (cyan fluorescent protein, CFP) and an acceptor domain (yellow fluorescent protein, YFP) as well as a bacterial ε-subunit of the F_0_F_1_-ATP synthase, which contains the ATP-binding site (Imamura et al., 2009). The distance and orientation of the donor and the acceptor fluophores and, therefore, the subsequent FRET depend on the conformation of the ε-subunit of the F_0_F_1_-ATP synthase, which is dependent on the ATP concentration (K_D_=1.2 mM) (Imamura et al., 2009). The ratio of the background-corrected signal from the CFP and YFP was the semi-quantitative readout for the intracellular ATP concentration ([ATP]_i_). Thus, percentual signal changes are not strictly equivalent to changes in [ATP]_i_.

Fluorescence was low pass filtered (690 nm) and split by a dichroic mirror (D510 nm) to capture CFP and YFP fluorescence separately. CFP emission was further bandpass filtered (median wavelength 480 nm/ bandwidth 40 nm, F47-480SG, AHF Analysentechnik AG, Tübingen, Germany).

#### Experimental protocols

All experiments began with baseline imaging (5-10 stacks, duration of at least 2.5 minutes) before KCl-induction of SDs or OGD. We investigated SDs in three conditions: 1) KCl-induced SDs in presence of oxygen and glucose, 2) SDs occurring during continuous OGD, and 3) SDs occurring during OGD with re-supply of oxygen and glucose following visual SD detection on LFP and [K^+^]_o_-recordings.

#### Data analysis

All data were analyzed in Igor Pro 8 (WaveMetrics, Inc, Lake Oswego, Oregon, USA) and Matlab (The MathWorks, Natick, Ma, USA, version R2022a). SDs were visually identified by their characteristic signatures in IOS (post-hoc analysis) as well as LFP and [K^+^]_o_ recordings (post-hoc and live analysis), respectively. The IOS during SDs was compared to the baseline IOS (prior to KCl-induced SDs or OGD) by calculating the percentage change.

Nernst potentials recorded with K^+^-sensitive electrodes were converted to [K^+^]_o_ in mM using the Nernst equation and assuming baseline [K^+^]_o_ to be equivalent to the K^+^ concentration in the aCSF (Schoknecht et al., 2017).

The ratio of YFP and CFP fluorescence (YFP/CFP) emitted by the ATP-sensor was used as a semi-quantitative parameter for dynamic ATP analysis. As a pre-processing step background signals were subtracted, e.g. derived from sensor-free extracellular compartments in high magnification images or from lateral ventricles in low magnification images. Further, z-planes were compressed into maximum intensity z-projections. Movement in x and y direction was corrected using a registration algorithm based on Fast Fourier Transformation (Guizar-Sicairos et al., 2008). To extract temporal ATP profiles, ROIs of individual cells, cortical and subcortical gray matter and white matter were manually selected. ATP profiles were self-normalized to their baseline average prior to OGD or SD induction.

#### Data reporting and statistical analysis

This is an exploratory study. Due to the lack of prior evidence sample sizes could not be accurately estimated and were selected to comply with likewise research. Slices were excluded from analysis if movement artifacts prevented reliable analysis. Neurons that moved out of the vertical stacks during the acquired time series were excluded from analysis. Neurons with low baseline YFP/CFP ratios, similar to those cells that had undergone SD under OGD, were also excluded.

Data are reported as median and interquartile 1 (IQ1, 25^th^ percentile) and IQ3 (75^th^ percentile). Boxes in boxplots display median, and IQ1 and IQ3 mark lower and upper box limits. Whiskers extend to minimal and maximal values unless considered as outliers (‘+’, values >1.5*IQR). Data analysis was not blinded. Statistical inference was performed by Wilcoxon test for paired comparisons and Mann-Whitney U-test for independent sample. Multiple comparisons were corrected by the Bonferroni post-hoc test. Differences were considered significant at p<0.05. Data will be made available upon reasonable request.

## Acknowledgement

We would like to thank Gudrun Bethge and Doris Grieshammer for their technical assistance, Dr. Agustin Liotta for his support with the pO_2_ recordings, and Marie-Elisabeth Burkart for her support in designing the figures.

---

**Supplementary Fig. 1 Control experiments**

A. Average of ATeam signals from 17 cortical neurons monitored for >50 minutes (imaging interval of 2 min) without experimental interventions, self-normalized to the mean of the monitoring period.
B. Maximum intensity z-projection before (left), during (middle), and after (right) perfusion of 30% hypoosmolar aCSF showing ATeam signals of the same individual cells displayed in Fig. 2Da. Note how the perfusion of hypoosmolar aCSF induces neuronal swelling but no reduction in [ATP]_i_.
C. Summary plot of ATeam signals from 20 cortical neurons (4 slices, 2 mice) during perfusion of 30% hypoosmolar aCSF.

