## Supplementary figures and images for "Spreading depolarizations exhaust neuronal ATP in a model of cerebral ischemia"

### Supplementary Fig. 1

**Supplementary Fig. 1.**

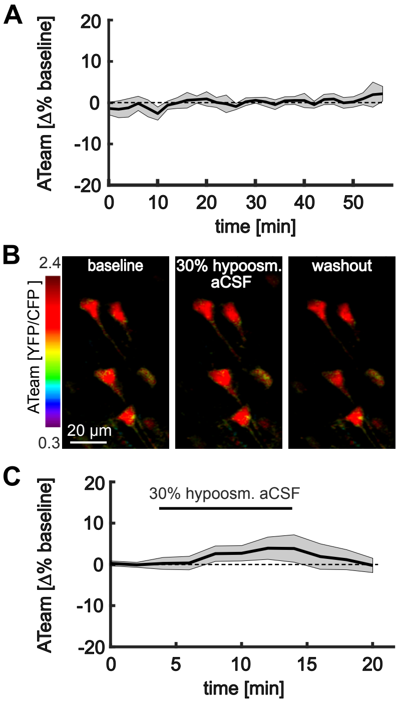
